# Marine snow as a habitat for microbial mercury methylators in the Baltic Sea

**DOI:** 10.1101/2020.03.04.975987

**Authors:** Eric Capo, Andrea Garcia Bravo, Anne L. Soerensen, Stefan Bertilsson, Jarone Pinhassi, Caiyan Feng, Anders F. Andersson, Moritz Buck, Erik Björn

**Affiliations:** Department of Chemistry, Umeå University, Umeå, Sweden; Department of Marine Biology and Oceanography, Institute of Marine Sciences, Spanish National Research Council (CSIC), Barcelona, Spain; Department of Environmental Research and Monitoring, Swedish Museum of Natural History, Stockholm, Sweden; Department of Aquatic Sciences and Assessment, SLU Uppsala, Sweden; Centre for Ecology and Evolution in Microbial Model Systems - EEMiS, Linnaeus University, Kalmar, Sweden; Department of Gene Technology, KTH Royal Institute of Technology, Science for Life Laboratory, Solna, Sweden

## Abstract

Methylmercury (MeHg), a neurotoxic compound biomagnifying in aquatic food webs, can be a threat to human health via fish consumption. However, the composition and distribution of the microbial communities mediating the methylation of mercury (Hg) to MeHg in marine systems remain largely unknown. In order to fill this gap of knowledge, we used the Baltic Sea Reference Metagenome (BARM) dataset to study the distribution of the genes involved in Hg methylation (the *hgcAB* gene cluster). We determined the relative abundance of the *hgcAB* genes and their taxonomic identity in 81 brackish metagenomes that cover spatial, seasonal and redox variability in the Baltic Sea water. The *hgcAB* genes were predominantly detected in anoxic water, but some *hgcAB* genes were also detected in hypoxic and normoxic waters. Higher relative quantities of *hgcAB* genes were found in metagenomes from marine snow compared to free-living communities in anoxic water, suggesting that marine snow are hotspot habitats for Hg methylators in oxygen-depleted seawater. Phylogenetic analysis identified well-characterized Hg methylators such as Deltaproteobacteria in oxygen-depleted water, but also uncovered Hg methylators within the Spirochaetes and Lentisphaerae phyla. Altogether, our work unveils the diversity of the microorganisms mediating MeHg production in the Baltic Sea and pinpoint the ecological niches of these microorganisms within the marine water column.

## Introduction

Methylmercury (MeHg) is a neurotoxic compound that accumulates in aquatic food webs and may be a threat to human health related to fish consumption (Mason *et al.*, 2012). Methylation of inorganic mercury (Hg) to MeHg is predominantly a biological process driven by anaerobic bacteria and archaea carrying the *hgcA* and *hgcB* genes (Parks *et al.*, 2013) and takes place in various oxygen-deficient environments (e.g. sediment, water, soil). Hg methylation appears to be controlled by the activity of Hg-methylating microbes, the composition and activity of microbial communities (that indirectly modulate Hg methylation), and Hg bioavailability (Bravo and Cosio, 2019). It is broadly established that the capacity for Hg-methylation is limited to specific microbial lineages, with the most commonly reported groups found in the Deltaproteobacteria and Methanomicrobiales (Gilmour *et al.*, 2013; Podar *et al.*, 2015; Bravo *et al.*, 2018; Yu *et al.*, 2018). However, a recent work has unravelled a higher phylogenetic diversity of microbes carrying the *hgcAB* genes than previously expected (McDaniel *et al.*, 2020) and this calls for novel analyses of microbial Hg-methylation in aquatic environments.

Recent advances in metagenomics have yielded new insights into the microbial taxonomic and functional diversity in various aquatic ecosystems (e.g., Mehrshad *et al.*, 2016; Haro-Moreno *et al.*, 2018; Nowinski *et al.*, 2019). The approach has for example been applied to broadly assess the presence and diversity of genes central to biological Hg cycling in marine systems (Podar *et al.*, 2015; Gionfriddo *et al.*, 2016; Bowman *et al.*, 2019; Villar *et al.*, 2020). Podar et al. (2015) only detected *hgcAB* genes in a few metagenomes from marine pelagic waters (seven out of 138 metagenomes) but highlighted that limited sequencing depths of these metagenomes could have hampered detection. A more recent study did not detect *hgcA* genes in waters from the Arctic and equatorial Pacific Oceans (Bowman *et al.*, 2019). Interestingly, the presence of *hgcAB*-like genes was reported in normoxic water from open ocean and sea ice in Antarctica, with a fraction of those genes being associated to microaerophilic nitrite oxidizing bacteria (Gionfriddo *et al.*, 2016; Villar *et al.*, 2020). Further, Blum et al. (2013) demonstrated that between 20 and 40 % of MeHg measured in surface mixed layer of the North Pacific Ocean originated from internal production in the surface water.

Marine snow (organic-rich particulate matter and aggregates) is hypothesized to provide both substrates for heterotrophic microbes (Azam and Long, 2001; Azam and Malfatti, 2007) and various anaerobic microenvironments (Alldredge and Silver, 1988; Bianchi *et al.*, 2018) that could potentially favor Hg methylation, via e.g. microbial sulfate-reduction. Based on this, several studies proposed (Lehnherr *et al.*, 2001, Monperrus *et al.*, 2007, Cossa *et al.*, 2009, Sunderland *et al.*, 2009, Schartup *et al.*, 2015) or demonstrated (Oritz *et al.*, 2015; Gascón Diez *et al.*, 2016) Hg methylation in settling particles. For the Baltic Sea it has been proposed that Hg methylation in normoxic water can be associated with phytoplankton blooms via production of increased levels of phytoplankton derived OM (marine snow) sinking through the water column (Soerensen *et al.*, 2016) providing suitable anoxic niches for Hg methylators. However, as far as we know there are no studies on microbial communities in relation to this phenomenon in the Baltic Sea or elsewhere. In addition to the oxygen-deficient microzones in marine snow, Oceans and coastal seas, such as the West Coast of South America, the Arabian Sea and the Baltic Sea, have experienced increased deoxygenation since at least the middle of the 20^th^ century (Breitburg *et al.*, 2018). This phenomenon can be caused by (i) warming that decreases the solubility of oxygen in the ocean and (ii) nutrient enrichment of coastal water causing an increase of algal biomass and subsequent decomposition of sinking organic matter by microbes consuming the oxygen (Breitburg *et al.*, 2018). Such oxygen deficient waters potentially offer ecological niches suitable for Hg-methylating microorganisms. Overall there are still important knowledge gaps concerning the process of Hg methylation in aquatic systems, in particular regarding variable redox conditions.

The Baltic Sea is an ecosystem that has experienced large increases in nutrient loads and oxygen consumptions over the last century, resulting in extensive coastal and offshore zones with permanent hypoxic and anoxic water below the oxygenated surface water (Conley *et al.*, 2011; Carstensen *et al.*, 2014). As such, the Baltic Sea represents a model for the expansion of coastal ecosystems influenced by anoxia. Elevated MeHg concentrations in the Baltic Sea have been observed in anoxic water (> 1000 fM) compared to hypoxic and normoxic water (Kuss *et al.*, 2017; Soerensen *et al.*, 2018). Soerensen et al. (2018) demonstrated that this was caused by increased rates of Hg methylation in the oxygen deficient water zones. They hypothesized that this process is predominantly driven by microbial sulfate-reduction because of the relatively high concentrations in dissolved sulfide in the anoxic water (up to ∼60 µM S^-II^). Although the concentrations of MeHg in normoxic water were generally low (13-80 fM), concentrations were higher than what could be explained by MeHg input form external sources only, and the authors inferred *in situ* formation of MeHg at a low rate also in normoxic water zones in the Baltic Sea (Soerensen *et al.*, 2016, 2018). The presence and distribution of microorganisms carrying the *hgcAB* genes, including taxa known to reduce sulfate, could unequivocally confirm the potential for *in situ* MeHg formation the Baltic Sea.

In this study, we assessed the spatial and seasonal variability of the *hgcAB* in the Baltic Sea, including water column profiles, allowing us to investigate for the first time the presence and variation of Hg-methylating microbes across redoxclines in the Baltic Sea. We revealed the presence and relative abundance of Hg methylators in both hypoxic and anoxic water masses of the Baltic Sea while *hgcAB genes* were present in very low abundance or not at all detected in normoxic waters. In addition, we found a higher proportion of *hgcAB* DNA sequences in metagenomes obtained from 3.0 µm filters compared to 0.2 µm filters, suggesting marine snow as an important habitat for Hg methylators. Our work provides new information on Hg-methylating microorganisms in coastal seas impacted by oxygen deficiency.

## Material and Methods

The Baltic Sea Reference Metagenome (BARM) data (Alneberg *et al.*, 2018) used in our study is composed of 81 metagenomes combined from three datasets spanning 13 locations (Figure 1) and selected to cover natural variation in geography, depth and seasons of Baltic Sea waters. A summary on sampling and filtration of water samples for each of the datasets is provided in Table 1. We classified the water samples into three categories based on the measured O_2_ concentrations: (i) normoxic water with O_2_ concentrations exceeding or equal to 2 mL O_2_.L^-1^ (ii) hypoxic water with detectable O_2_ concentrations lower than 2 mL O_2_.L^-1^ and (iii) anoxic water with no detectable O_2_. Methods for DNA extractions, library preparations and sequencing as well as the initial processing of metagenomics data is described in greater details in Alneberg et al. (2018). Briefly, preprocessed reads were co-assembled using Megahit (version 1.0.2, Li et al 2015). Functional and taxonomic annotation was performed for the 6.8 million genes found on the 2.4 million contigs >1 kilobase. In order to detect *hgcAB* homologs genes, we first analyzed the 6.8 million predicted genes with the function hmmsearch from the hmmer software (3.2.1 version, Finn *et al.*, 2011) with the use of the HMM profiles of *hgcAB* protein sequences from Podar et al (2015) as a reference database (Supplementary Text S1). We considered genes with E-values ≤ 10^−3^ as significant hits resulting in a total of 3,215 genes. Only a fraction of these 3,215 genes correspond to *hgcA* and *hgcB* genes and we therefore performed a manual check of their protein sequences using the knowledge from the seminal paper of Parks et al. (2013) that described unique motifs in *hgcA* (NVWCA(A/G/S)GK) and *hgcB* protein sequences (C(M/I)EC(G/S)(A/G)C). For phylogenetic analysis, we used the protein sequences of the 650 *hgcAB* gene clusters identified as putative Hg methylators by McDaniel et al. (2020). These 650 *hgcAB* gene clusters were obtained from the analysis of publicly available isolate genomes and metagenome-assembled genomes (Supplementary Text S2, McDaniel *et al.*, 2020). For each phylogenetic analysis, protein sequences were aligned using MUSCLE (cluster method UPGMA) in the software MEGAX (Kumar *et al.*, 2018) and approximate maximum likelihood (ML) trees were constructed using FastTree (Price *et al.*, 2009). The trees were visualized using iTOL (Letunic and Bork, 2019) and clades were collapsed by the dominant, monophyletic phyla when possible for visualization ease.

**Table 1.**
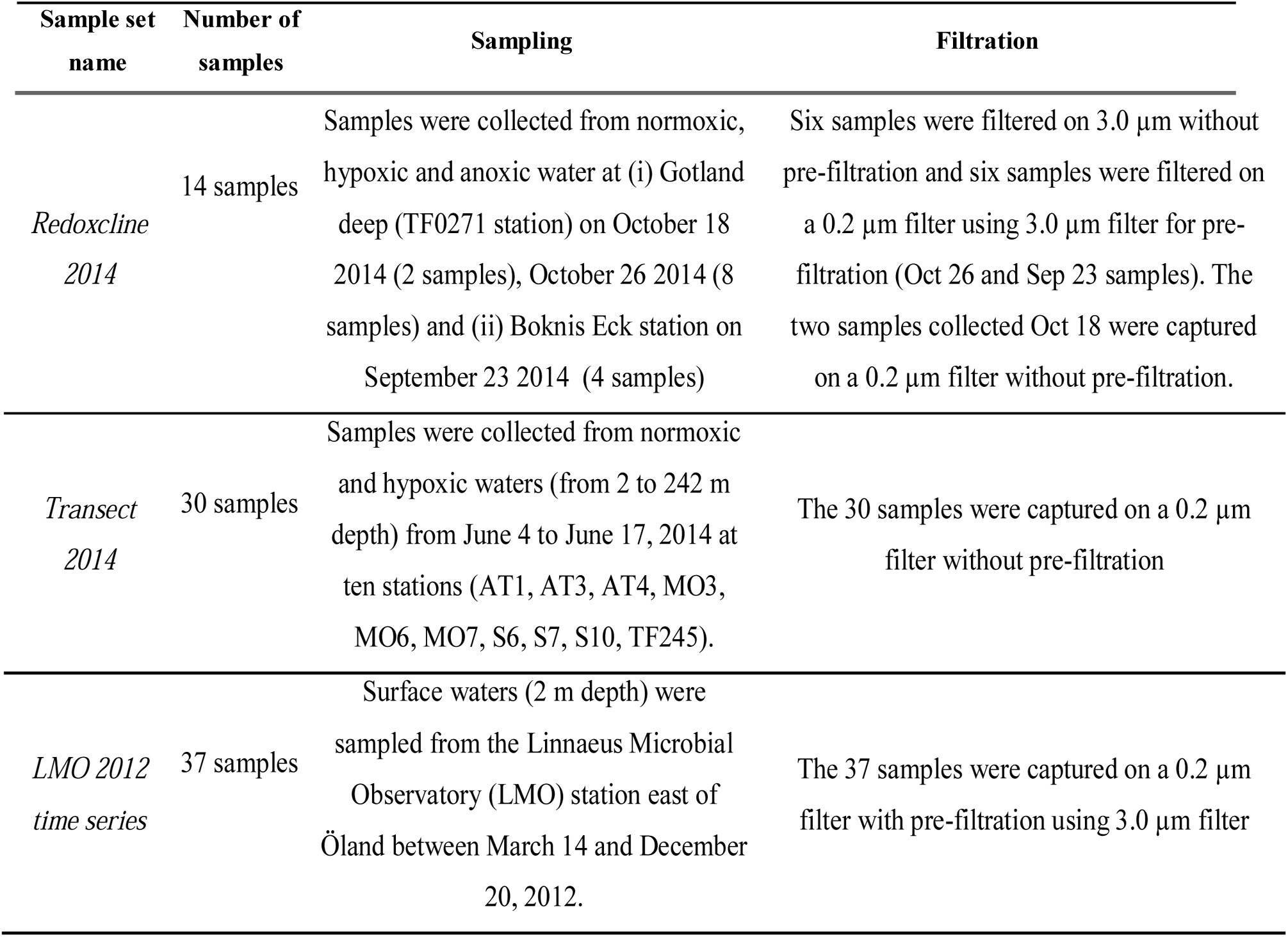
Description of the number of metagenomes obtained with their respective sampling and filtration strategies for each of the three datasets.

**Figure 1.**
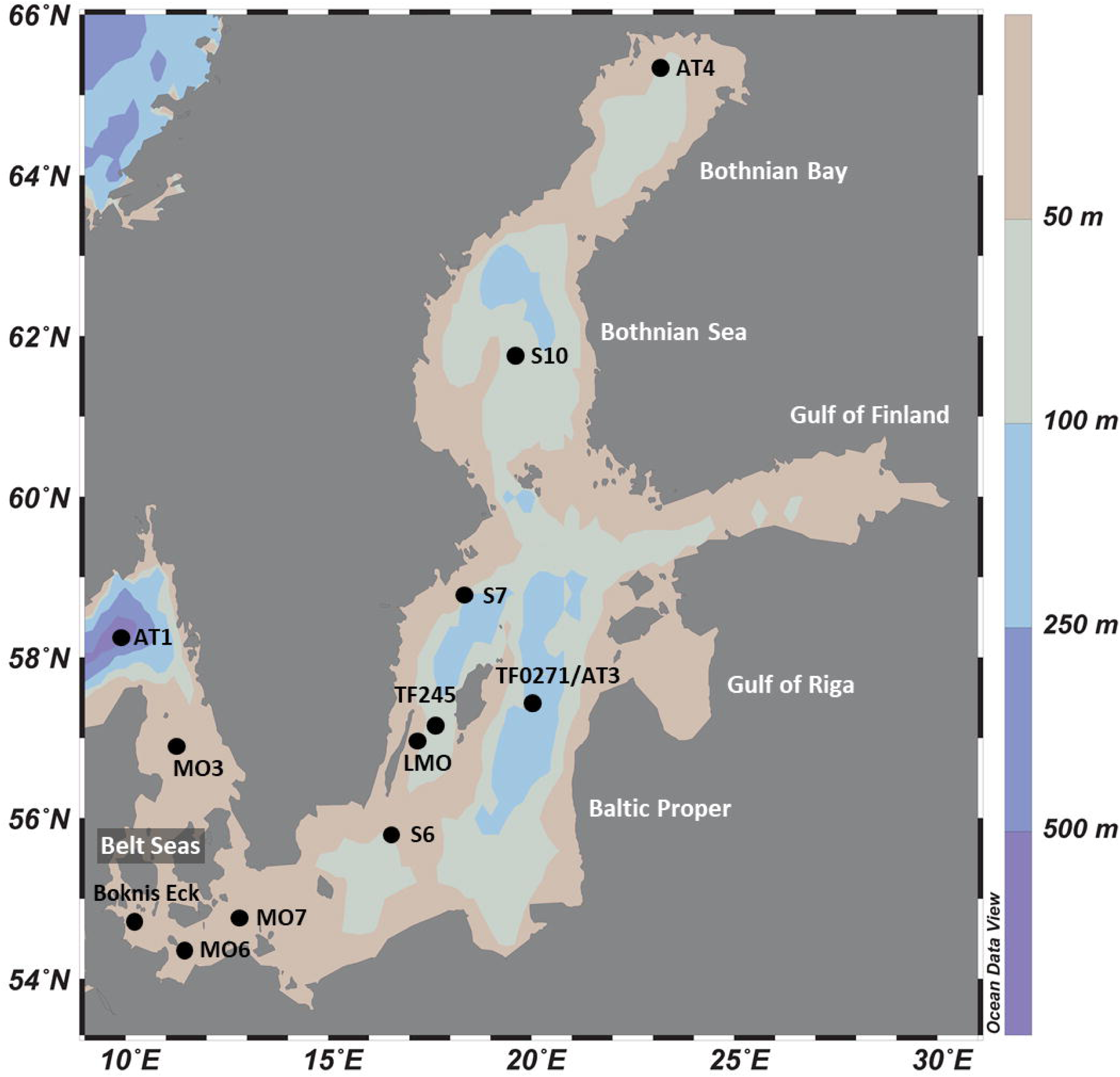
Location of study sites in the Baltic Sea

## Results

### Detection of *hgcAB* genes across the Baltic Sea metagenomes

The total DNA sequence counts (number of reads) in the 81 Baltic Sea metagenomes ranged from 1.6 to and 37 million reads (mean 12.9, sd: 9.7). Among the total of 6.8 million protein-coding genes predicted from the co-assembly, 22 *hgcA*-like and 12 *hgcB*-like genes were detected. In some cases, *hgcA* and *hgcB* genes were found side-by-side on the same contig. Overall, we detected: (i) nine *hgcAB*-like gene clusters, (ii) 13 *hgcA*-like genes, and (iii) three *hgcB*-like genes (Figure 2, Supplementary Text S3). The resulting 25 gene clusters or single genes were named as displayed in Figure 2. The thirteen *hgcA*-like genes found without a coupled *hgcB* gene were always found at an extremity of the respective contig, possibly explaining why no *hgcB* genes were detected alongside. In contrast, the three *hgcB*-like genes found alone on their respective contig were consistently found in the central portions of the contigs, with no downstream or upstream protein motifs from *hgcA* genes. Most of the *hgcA* and *hgcB* genes had the most common protein motifs, NVWCAAGK and CMECGAC, respectively, as described by Parks et al. (2013). Two gene clusters contained both the “NVWCASGK” and “CIECGAC” motifs (BARM-01 & -09) while BARM-07 was the only gene cluster with the NVWCAAGK and CIECGAC combination (Figure 2).

**Figure 2.**
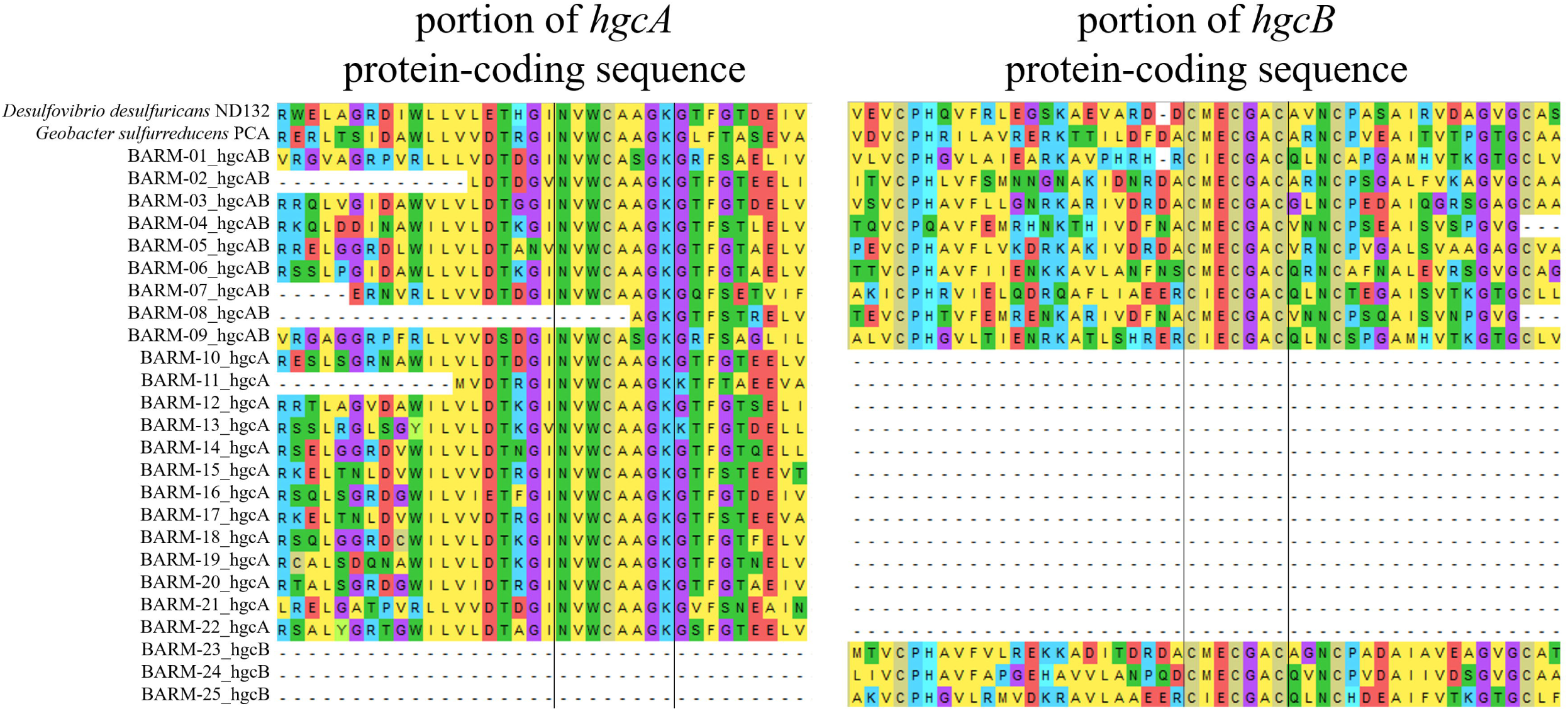
Alignment of conserved regions for protein sequences of the nine hgcAB-like gene clusters, (BARM-01 to 09) 13 hgcA-like genes (BARM-10 to 22) three hgcB-like genes (BARM-23 to 25) detected in the BARM dataset. The corresponding hgcAB Protein sequences from Desulfovibrio desulfuricans ND132 and Geobacter sulfurreducens PCA were added in the alignment as references.

### Taxonomic identifications of *hgcAB*-like genes found in the Baltic Sea

In order to taxonomically identify each Hg methylation gene detected in the BARM dataset, we constructed phylogenetic trees using the 650 *hgcAB* gene clusters generated by McDaniel et al. (2020). For the nine *hgcAB*-like gene clusters concatenated, we performed a phylogenetic analysis with the 650 *hgcAB* gene clusters (Figure 3, Supplementary File S1). For the 13 *hgcA* and three *hgcB* genes detected alone in their respective contig, two additional phylogenetic trees were performed using the 650 *hgcA* and 650 *hgcB* genes, respectively (Supplementary File S2 and S3). The phylogenetic analysis revealed the presence of several *hgc* genes (referring hereafter either to *hgcAB* gene clusters, *hgcA* genes or *hgcB* genes) affiliated with the order Desulfobacterales (Deltaproteobacteria, class Desulfobacterota in GTDB classification): a member of the family Desulfobulbaceae (BARM-15 & -17), a *Desulfobacula* sp. (BARM-04 & -08), and a *Desulforhopalus* sp. (BARM-11) (Figure 3). In addition, the *hgc* genes BARM-02, -06 & -10 were associated with members of the orders Desulfatiglanales (naphS2 family) and Desulfarculales (Desulfarculaceae family) and Syntrophales. The hgc genes BARM-01, -07, -09 & -21 clustered together and were closely related to genes detected in various microbial phyla, with the closest related *hgc* genes detected in the genomes of two Spirochaetes from the family Treponemataceae. Three hgc genes (BARM-05, -19 & -22) were associated with members of the Lentisphaerae phylum, which is part of the widespread PVC superphylum (i.e. including Planctomycetes, Verrucomicrobia, Chlamydiae and Lentisphaerae). In addition, some BARM *hgc* genes were closely related to Firmicutes (Clostridia, BARM-14) and a group of *hgc* genes from Euryarchaea and Chloroflexi clustered together (BARM-18). Finally, seven BARM hgc genes were associated with clades including various microbial lineages and are thus classified here as unidentified (BARM-03, -12, -13, -16, -20, -23 & -24).

**Figure 3.**
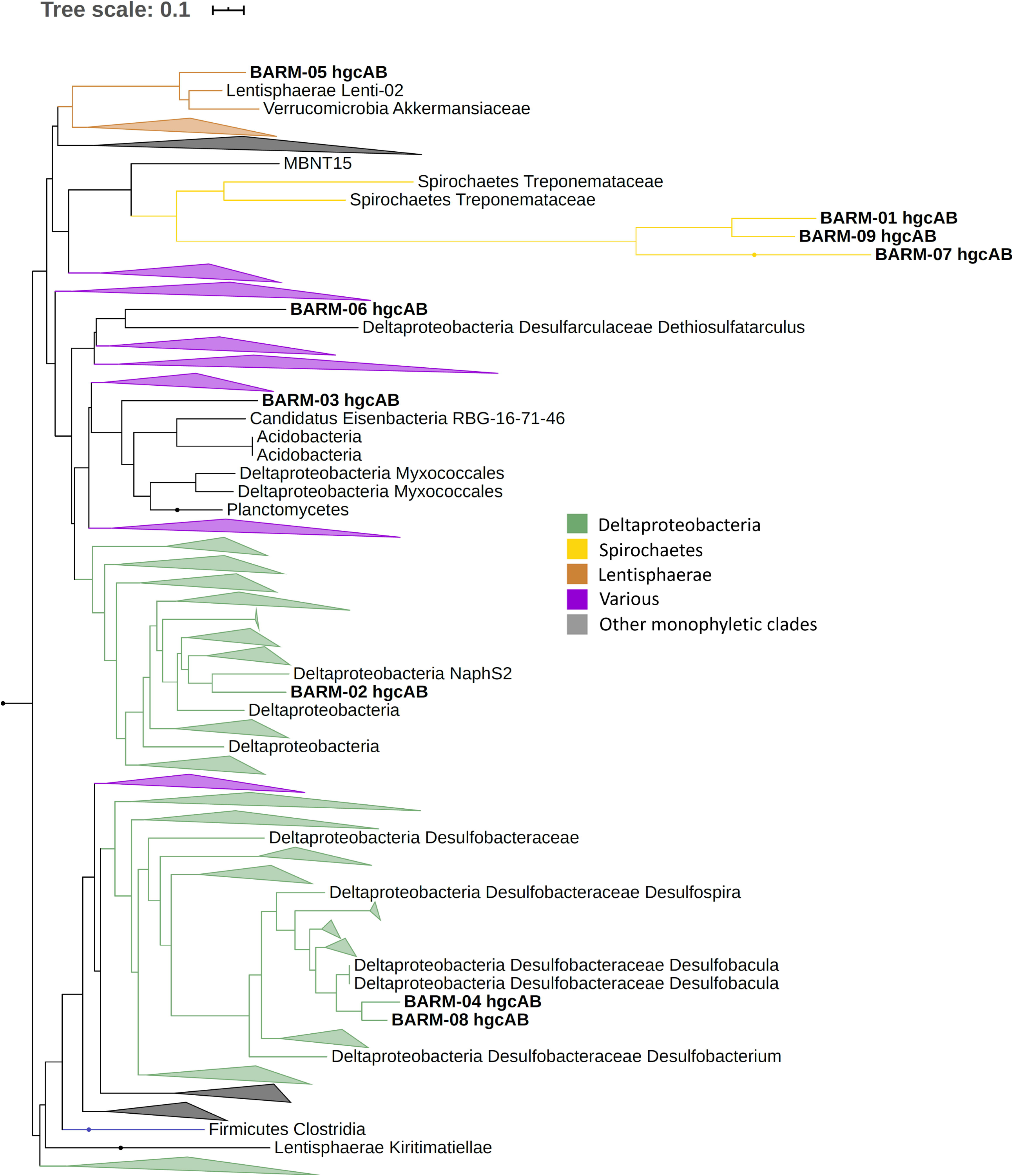
Phylogenetic tree of hgcAB gene clusters with collapsed clades. Collapsed clades are discriminated with colors based on their taxonomy. Gene clusters detected in the Baltic Sea water column (BARM) are displayed in bold.

### Relative abundance of *hgcAB* genes in the Baltic Sea water column

The summed relative abundance of *hgcA* and *hgcB*-like genes in the Baltic Sea water column (i.e. the number of reads for the *hgcAB*-like genes per total annotated reads per sample, expressed as %) ranged from undetected to 6.7 × 10^−3^ % (mean: 0.3 × 10^−3^, sd: 1.2 × 10^−3^) (Table 2, Figure 4). The highest relative abundance of *hgcAB*-like genes was found in the hypoxic water from 76.5 m depth at station S7 with 6.7 × 10^−3^ % (Figure 4). Elevated abundance of *hgcAB*-like genes was also found in hypoxic and anoxic water from the TF0271/AT3 station with the highest values at a water depth of 200 m with 6.5 × 10^−3^ %. The proportion of *hgcAB* genes detected in the other 10 locations was relatively low with a maximum value of 0.2 × 10^−3^ % in the hypoxic layer (87.5 m) at station TF245 (Figure 4). At the LMO station, for which only normoxic water was sampled, the highest proportion of *hgcAB* genes found in the 37 samples was less than 0.01 × 10^−3^ % (sample LMO 120806, Supplementary Table S1). We found that the most abundant *hgcAB* genes in Baltic Sea anoxic water belonged to members of Deltaprotebacteria, more specifically members of Desulfobulbaceae, Desulfarculaceae, *Desulfobacula* sp. and Syntrophales, and Spirochaetes from the Treponemataceae family (Figure 4, Table S3).

**Table 2:**
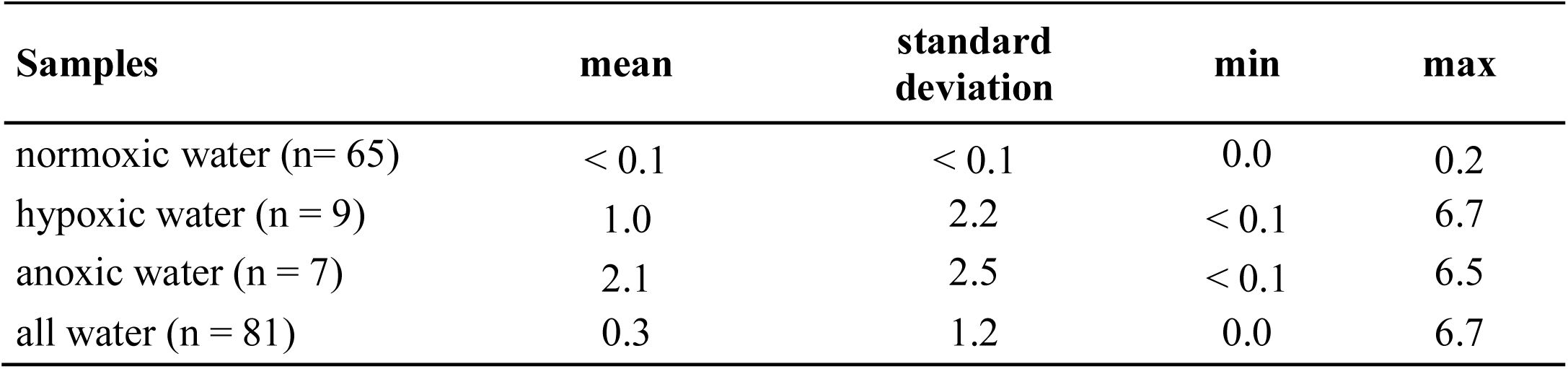
Proportion (units in 1 × 10^−3^; normalized to total DNA sequences count of each metagenome) of *hgcAB*-like genes in the 81 metagenomes. The values obtained from the metagenomes collected in the three type of water defined in this study (normoxic, hypoxic and anoxic water) are displayed in this table.

**Figure 4.**
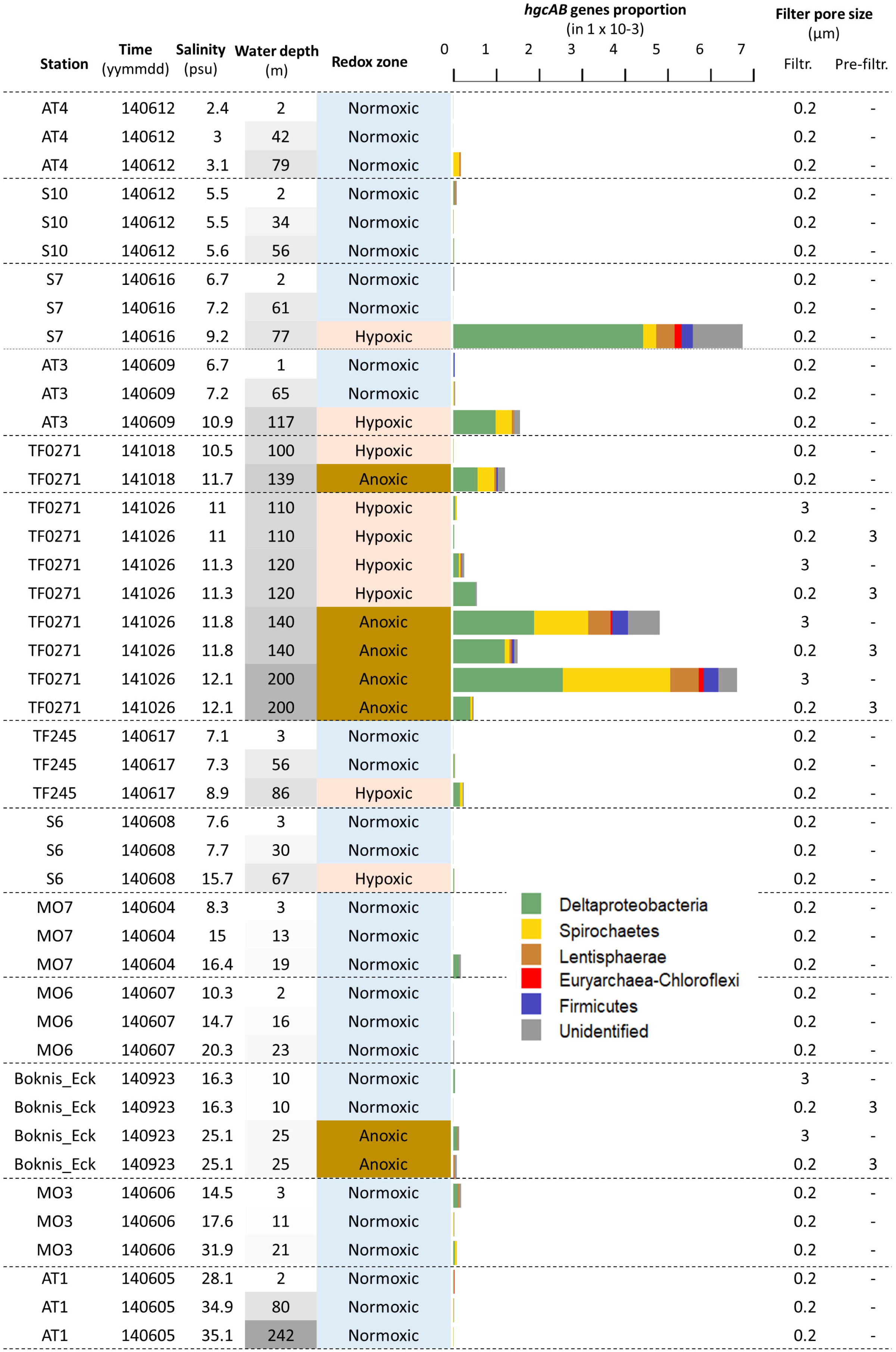
Relative abundance of hgcA- and hgcB-like genes in samples from the dataset “Redoxcline 2014” and “Transect 2014”. The dataset “LMO 2012” is not included because only few genes were detected and at low abundances (data shown in Supplementary Table S1). The sampling time is written using the YYMMDD format. The sampling depth (m) is color coded based on its respective gradient (darker shade of gray with increasing depth). The water redox zone is color coded based on the O2 categories defined in the text (light blue for normoxic, beige for hypoxic and brown for anoxic conditions). The abbreviations “Filtr.” and “Pre-filtr.” indicate the pore size of the filters used to obtain each metagenome and if pre-filtration were done prior to filtration, respectively.

### Differences in *hgcAB* genes relative abundance with filter-size

The quantity of *hgcAB*-like genes detected in metagenomes obtained from the Baltic Sea water column differed systematically between filter size fractions (Figure 4). Metagenomes obtained from the TF0271 station profile, and filtered onto 3.0 µm filters (hereafter referred to as the “3.0 µm metagenomes”), had consistently higher proportions of *hgcAB*-like genes (up to 6.5 × 10^−3^ %) than metagenomes obtained from 0.2 µm filters following pre-filtration with 3.0 µm filters (hereafter referred to as the “0.2-3.0 µm metagenomes”) (up to 1.5 × 10^−3^ %) (Figure 4). This was especially clear for the anoxic TF0271-Oct26 samples, where the relative abundance of *hgcAB* genes was three and 14 times higher in 3.0 µm metagenomes compared to 0.2-3.0 µm metagenomes for samples collected at 140 and 200 m depth, respectively (Figure 4, Supplementary Table S1). At Boknis Eck, only a few samples were collected and the gene proportions were too low to properly investigate differences between filtration methods. At TF0271, *hgcAB* genes from Deltaproteobacteria were predominant for both 0.2-3.0 and 3.0 µm metagenomes (Figure 4). In these metagenomes, the most abundant *hgcAB* genes were attributed to Deltaproteobacteria from the families Desulfobulbaceae and Desulfarculales (Supplementary Table S1). The 3.0 µm metagenomes from anoxic TF0271 samples (both 140 and 200 m) featured higher proportions of genes affiliated with Spirochaetes and Lentisphaerae (Figure 4).

## Discussion

Our phylogenetic analysis of the Baltic Sea Reference Metagenome dataset (Alneberg *et al.*, 2018) identified at least 18 different *hgcAB* gene clusters that belong to several microbial lineages (Figure 3, Supplementary File S2 and S3). The majority of *hgcAB* genes detected were affiliated with Deltaproteobacteria (or Desulfobacterota with GTDB classification) notably in genomes from sulfate reducing bacteria (e.g., Compeau and Bartha, 1984, 1985, Gilmour *et al.*, 2013). Some of the identified Deltaproteobacteria/Desulfobacterota belong to groups of organisms previously known or predicted to perform Hg methylation, such as a species of *Desulfobacula* (Gionfriddo *et al.*, 2016), members of the Desulfobulbaceae family (Benoit *et al.*, 2001, Colin *et al.*, 2018), and a member of the naphS2 group (Parks *et al.*, 2013). It is noteworthy that none of the *hgcAB* genes detected in the water of the Baltic Sea were closely related to iron reducing bacteria such as the Geobacteraceae family (Fleming *et al.*, 2006; Kerin *et al.*, 2006; Bravo *et al.*, 2018). However, relatively high Fe(II) concentrations were previously reported in anoxic water from the Baltic Sea (> 25 nM; Pohl and Fernandez-Otero, 2012) but with seasonal and inter-annual variability in concentrations. Our finding that none of the *hgcAB* was closely related to Geobacteraceae suggests that other iron reducing bacteria without the capability to methylate Hg, such as *Shewenella* sp. (incl *S. baltica*), are responsible for the formation of Fe(II) in the Baltic sea water column. Indeed, Shewanellaceae are in general one to two orders of magnitude more abundant than Geobacteraceae in the metagenomes compiled in the BalticMicrobeDB (https://barm.scilifelab.se). Our results thus suggest that Hg methylation is not linked to iron-reduction in the Baltic Sea. Microbial syntrophy, defined as an obligate mutualistic metabolism, is a process known to occur mainly in environments with shortage of favorable electron acceptors, e.g. mostly in anoxic environments (McInerney *et al.*, 2009; Morris *et al.*, 2013). Some syntrophic microbes were recently suggested to be involved in the Hg methylation process (Gilmour *et al.*, 2013; Yu *et al.*, 2018; Hu *et al.*, 2020), but we did not identify *hgcAB* genes from known syntrophic microorganisms in the Baltic Sea. Instead, we identified *hgcAB*-carrying bacteria in various groups that are still poorly described (McDaniel *et al.*, 2020) including members of Spirochaetes and Lentisphaerae phyla. For instance, among the predominant *hgcAB*-like genes detected in Baltic Sea, the *hgcAB* genes BARM-01, -07 & -09 were clustered together and found closely related to Spirochaetes (Figure 3). However, it should be noticed that these *hgcAB* genes were also grouped together with various other bacterial groups in the phylogenetic analysis, including MBNT15, Acidobacteria, Actinobacteria and Verrucomicrobia (Supplementary File S1 for full description), which precludes a more precise identification at present.

The 81 metagenomes collected from three sampling efforts (Alneberg *et al.*, 2018) cover substantial variations in water depth, season and location across the Baltic Sea and our study investigates the relationship between putative Hg methylating populations and these factors. We found higher relative abundance of *hgcAB* genes in oxygen deficient water (hypoxic and anoxic layers) compared to those observed in normoxic layers also when including an extensive time series (37 time points over the year 2012) at 2 m depth. This finding is in agreement with the general understanding that methylation of Hg in aquatic systems is associated with anoxic conditions (Eckley *et al.*, 2005, Malcolm *et al.*, 2010; Compeau and Bartha, 1984, Gilmour *et al.*, 1992, Bravo *et al.*, 2015). Several studies (Goñi-Urriza et al., 2015; Bravo et al., 2016, Christensen et al., 2019) have however demonstrated a poor quantitative relation between Hg methylation rate and the presence of *hgcA* genes (mRNA and DNA), likely because of additional important factors/processes affecting Hg methylation. Indeed, while Soerensen et al. (2018) found Hg methylation rate constants below detection limit in both hypoxic and normoxic waters of the Baltic Sea, we find a relatively high abundance of *hgcAB*-like genes in hypoxic water samples at two locations (AT3, 117 m and S7 77m). This finding reveals the potential for Hg methylation in hypoxic waters.

*In situ* formation of MeHg in normoxic waters has been proposed to occur in anaerobic microzones formed around organic-rich particulate matter and aggregates, also called marine snow, by several studies (Lehnherr *et al.*, 2001, Monperrus *et al.*, 2007, Cossa *et al.*, 2009, Sunderland *et al.*, 2009, Schartup *et al.*, 2015) and experimentally demonstrated in a few cases (Oritz *et al.*, 2015; Gascon Diez *et al.*, 2016). Our study expands on the role of marine snow for Hg methylation in coastal seas and demonstrates that marine snow is an important habitat for Hg methylators also in anoxic water. Firstly, we found higher relative abundance of *hgcAB* genes in the metagenome of marine snow from anoxic (3.0 µm filters) compared to hypoxic (3.0 µm filters) or normoxic (not pre-filter with 3.0 µm filters) water samples (Figure 4). Considering that the 3.0 µm filters represent the particle and aggregated organic matter fraction (that in turn is frequently referred to as marine snow), our results suggest that marine snow becomes a more suitable habitat for Hg methylators (and thus potentially constitutes an environment with high Hg methylation rate) when reaching anoxic water. This phenomenon is likely caused by an increased prevalence of anaerobic conditions in the marine snow in anoxic water layers. Secondly, we found a higher proportion of DNA sequences of *hgcAB* genes in 3.0 µm metagenomes than in the free-living microbes, represented by the metagenomes obtained from 0.2-3.0 µm size fraction, in anoxic waters. This finding could be explained by the marine snow containing high concentrations of organic compounds suitable as metabolic electron donors for many microbial activities occurring in anoxic conditions (Bianchi *et al.*, 2018). It is noticeable that previous studies demonstrated differences between particle-associated and free-living bacterial communities in the Baltic Sea water column but that Spirochaetes and Lentisphaerae, for which *hgcAB*-like genes were found relatively abundant in the 3.0 µm metagenomes from BARM dataset, were not identified as more predominant in particle-associated bacterial communities (Rieck *et al.*, 2015, BalticMicrobeDB https://barm.scilifelab.se).

Our findings advance the understanding of the diversity and distribution of genes involved in Hg methylation as well the *hgcAB*-carrying microbial populations in marine environments. Particularly noteworthy was the finding that most of *hgcAB*-carrying microbes in the Baltic Sea water column were Deltaproteobacteria/Desulfobacterota predominantly found in oxygen-deficient zones (anoxic but also hypoxic zones). In addition, the differences in the relative abundance of *hgcAB* genes in metagenomes obtained from 0.2 compared to 3.0 µm pore size filters suggest marine snow as a niche for Hg methylating microbial communities and therefore and important potential hotspot for MeHg formation in both hypoxic and anoxic sea water zones. Finally, our phylogenetic analysis highlights that a substantial part of the Hg methylators present in the Baltic Sea, and likely in other marine environments, are still poorly described and more works are needed to isolate, characterize and describe their genetic diversity.

## Acknowledgements

This work was funded by the Swedish research council Formas (grant 2018-01031). The computations were performed on resources provided by SNIC through Uppsala Multidisciplinary Center for Advanced Computational Science (UPPMAX) under Project SNIC 2019/8-322.

## Captions of supplementary material

**Supplementary Table S1.** Proportion of *hgcAB* genes (out of the total number of reads of each metagenome) in each metagenome from the three datasets used in the present study. The protein sequences, taxonomic identifications associated to each gene are displayed. The environmental conditions associated to the water samples for each metagenome are displayed.

**Supplementary Text S1.** *HgcAB* protein sequences of HMM profiles modified from Podar et al. (2015)

**Supplementary Text S2.** Protein sequences of *hgcAB* gene clusters identified in 650 bacterial and archaeal genomes (McDaniel *et al.*, 2020).

**Supplementary Text S3.** Protein sequences of *hgcAB*-like genes detected in BARM dataset

**Supplementary File S1.** Extended phylogenetic tree of *hgcAB* genes performed from the protein sequences of 650 *hgcAB* gene clusters (McDaniel *et al.*, 2020) and BARM *hgcAB* genes (in newick format).

**Supplementary File S2.** Extended phylogenetic tree of *hgcA* genes performed from the protein sequences of 650 *hgcA* genes (McDaniel *et al.*, 2020) and BARM *hgcA* genes (in newick format).

**Supplementary File S3.** Extended phylogenetic tree of *hgcB* genes performed from the protein sequences of 650 *hgcB* genes (McDaniel *et al.*, 2020) and BARM *hgcB* genes (in newick format).

